# 40 Hz Audiovisual Stimulation Improves Sustained Attention and Related Brain Oscillations

**DOI:** 10.1101/2025.08.25.671937

**Authors:** Matthew K. Attokaren, Lu Zhang, Sindhura Mettupalli, Annabelle C. Singer

## Abstract

Gamma oscillations (30-100 Hz) have long been theorized to play a key role in sensory processing and attention by coordinating neural firing across distributed neurons. Gamma oscillations can be generated internally by neural circuits during attention or exogenously by stimuli that turn on and off at gamma frequencies. However, it remains unknown if driving gamma activity via exogenous sensory stimulation affects attention. We tested the hypothesis that non-invasive audiovisual stimulation in the form of flashing lights and sounds (flicker) at 40 Hz improves attention in an attentional vigilance task and affects neural oscillations associated with attention. We recorded scalp EEG activity of healthy adults (n=62) during one hour of either 40 Hz audiovisual flicker, no flicker as control, or randomized flicker as sham stimulation, while subjects performed a psychomotor vigilance task. Participants exposed to 40 Hz flicker stimulation had better accuracy and faster reaction times than participants in the control groups. The 40 Hz group showed increased 40 Hz activity compared to the control groups in agreement with previous studies. Surprisingly, 40 Hz subjects had significantly lower delta power (2-4 Hz), which is associated with arousal, and higher functional connectivity in lower alpha (8-10 Hz), which is associated with attention processes. Furthermore, decreased delta power and increased lower alpha functional connectivity were correlated with better attention task performance. This study reveals how gamma audiovisual stimulation improves attention performance with potential implications for therapeutic interventions for attention disorders and cognitive enhancement.

## INTRODUCTION

Gamma stimulation, lights and sounds that turn on and off at 40 Hz, known as “flicker”, has recently gained interest as a potential non-invasive intervention for Alzheimer’s disease^1–6^. Multiple studies have shown that chronic gamma flicker over days or weeks reduces Alzheimer’s pathology, including amyloid beta, altered neuroimmune activity — such as microglia function and cytokine signaling — and enhanced glymphatic clearance in mice^2,5–8^. Recent and ongoing studies are determining whether gamma flicker has beneficial effects in human patients with Alzheimer’s disease, epilepsy, and other diseases^1,3,9^. While prior studies have focused on the effects of chronic gamma flicker in disease, little is known about gamma flicker’s acute influences on cognition in healthy adults. A key function of endogenous gamma oscillations is to facilitate attention processing^10–16^, but how acute gamma sensory flicker might affect attention is unclear. Indeed, whether sensory-induced gamma enhances, coexists with, or disrupts ongoing oscillations and related cognitive functions is a topic of debate^17,18^.

Gamma oscillations (which can include oscillatory activity from 30-100 Hz) have long been theorized to play a key role in sensory processing and attention by coordinating firing across distributed neurons^19^. Attention is thought to be facilitated by internally generated gamma oscillations because this activity coordinates neurons across brain regions to fire together on short timescales, thus enhancing communication between regions^12,20–22^. Specifically, neurons that respond to the attended stimulus exhibit increased synchronization in the gamma frequency range compared to other neurons, and this synchronized activity is more likely to drive downstream activity.^11^ While gamma oscillations observed during attention are internally generated by neural circuits, gamma frequency activity is also elicited by exogenous stimuli that turn on and off at gamma frequencies, like 40 Hz^10,23^. Current studies have debated whether exogenously driven gamma oscillations from rhythmic sensory stimuli have roles overlapping with those of gamma oscillations generated internally. Some theories posit that entrainment of neural activity to sensory inputs enhances representations of attended stimuli^24^. Some evidence suggests that rhythmic sensory inputs entrain endogenous oscillations while other evidence indicates that sensory-driven and endogenous oscillations separately co-exist^18,24–27^.

As a result, it is unclear if stimulating gamma activity via gamma sensory flicker would enhance or disrupt functions that are facilitated by endogenous gamma, including attention. Flashing lights and sounds like those used during flicker may enhance sensory entrainment or distract an individual from a cognitive task. Thus, we explored how acute gamma audiovisual flicker affects vigilance, or the ability to attend and respond to unpredictable stimuli over a prolonged period, during a psychomotor vigilance task. To elucidate how sensory stimulation affects multiple aspects of neural activity, we asked how gamma flicker during a vigilance task influences other oscillations involved in arousal and attention, such as delta (typically defined as 1-4 Hz) and alpha (8-13 Hz) oscillations^28–31^. Because oscillations facilitate communication across networks^32–34^, we asked if gamma flicker affected functional connectivity, or the temporal coordination of neural activity, to enhance computation in attention networks.

In this study, we tested the hypothesis that non-invasive sensory stimulation in the form of flashing lights and sounds at 40 Hz improves attentional vigilance, and modulates neural oscillations and functional connectivity associated with attention processing. We exposed 61 healthy young adults to 40 Hz audiovisual flicker or control conditions while they performed a vigilance task, and we recorded EEG activity to assess neural dynamics during stimulation and attention processing. One control group received sham stimulation: a random, non-periodic flicker stimulation that had the same average on-time as the 40 Hz group to control for the effects of a flashing stimulus. The other control group was exposed to constant light that was measured to be the same brightness as the 40 Hz flicker stimulation. We found that 40 Hz flicker enhanced vigilance task performance, including faster reaction times and improved accuracy. As expected, 40 Hz flicker increased power in the gamma-flicker response band (39-41 Hz). However, beyond this, we observed that 40 Hz-exposed subjects had several unexpected changes *outside* of the gamma band. Notably, 40 Hz flicker decreased delta power, and lower delta power correlated with better vigilance across all participants. Additionally, 40 Hz specifically increased functional connectivity within the lower alpha (8-10 Hz) band, a frequency range linked to attentional demands and sensory processing^28,35–37^. This enhanced lower alpha band functional connectivity was positively correlated with better vigilance task performance. These findings demonstrate that 40 Hz flicker enhanced vigilance and modulated neural activity beyond the gamma band, revealing broader and more complex effects than previously recognized. Indeed, these results show that the effects of 40 Hz flicker extend beyond the gamma range and engage activity associated with arousal and attention. More broadly, these results highlight novel effects of acute sensory flicker in healthy adults and suggest potential applications for cognitive enhancement and therapeutic interventions.

## METHODS

### Participants

The study was approved by the Institutional Review Board at the Georgia Institute of Technology. Participants (32 males, 30 females) between the ages of 18 to 40 were analyzed in this study (**Table 1**). Participants were screened for eligibility based on the inclusion criteria of right-handedness. Exclusion criteria were conditions or medications that would affect blood flow and/or neural processes, color blindness and/or uncorrected vision, unremovable metal in the form of things like piercings or implants of any kind, history of neurological or psychological disorders/diseases, and any other conditions that could potentially affect the EEG results.

Participants were also screened for any conditions that may necessitate exclusion from our EEG study for safety or technical reasons including being unable to sit for long periods of time or being unable to tolerate flashing lights.

All participants provided written and verbal consent prior to their session and were informed of any potential risks that could occur as a result of study procedures. Participants were compensated monetarily or with class credit for their participation in the study. Participants were randomly assigned to one of the three stimulation groups: 40 Hz, Random, and Constant Light (also called “Light”). We found no significant differences in the sample demographics between groups (**Table 1**). Prior to the study, participants were asked to ensure they kept to their normal routines to minimize changes in their daily practices that could influence their EEG session. Participants were surveyed immediately before and after their session to assess changes in drowsiness, dizziness, and boredom during the study.

### Flicker Exposure

Participants were separated into three groups: 40 Hz (n=21), Random (n=22), and Constant Light (n=19) Stimulation. To administer the different stimulation types, participants were seated in front of a computer monitor with a frame of light-emitting diodes (LEDs) around the border that turned on and off with millisecond precision (**Fig 1B**). For the 40 Hz group, during the 1-hour stimulation period, participants were exposed to LEDs and sound delivered via ear buds that turned on and off every 12.5 msec (50% duty cycle) to generate 40 Hz stimulation (**Fig. 1C**). For the Random group, the LEDs and audio were set to turn on and off at a non-periodic, variable rate that maintained the same average number of on-off transitions as the 40 Hz group. For the Constant Light group, the LEDs remained continuously on without modulation, and no audio was provided. Participants did not know if they were in a control or treatment group. They were not informed about different stimulation groups and all subjects were exposed to flickering stimuli over the course of the experiment, thus they were unlikely to deduce their group assignment even though the stimulus was detectable. Due to the flicker stimulus being visible, experimenters were not blind to group during data collection. All subjects were analyzed together using the same processing pipeline and thus experimenters were blind during analysis.

**Figure 1.**
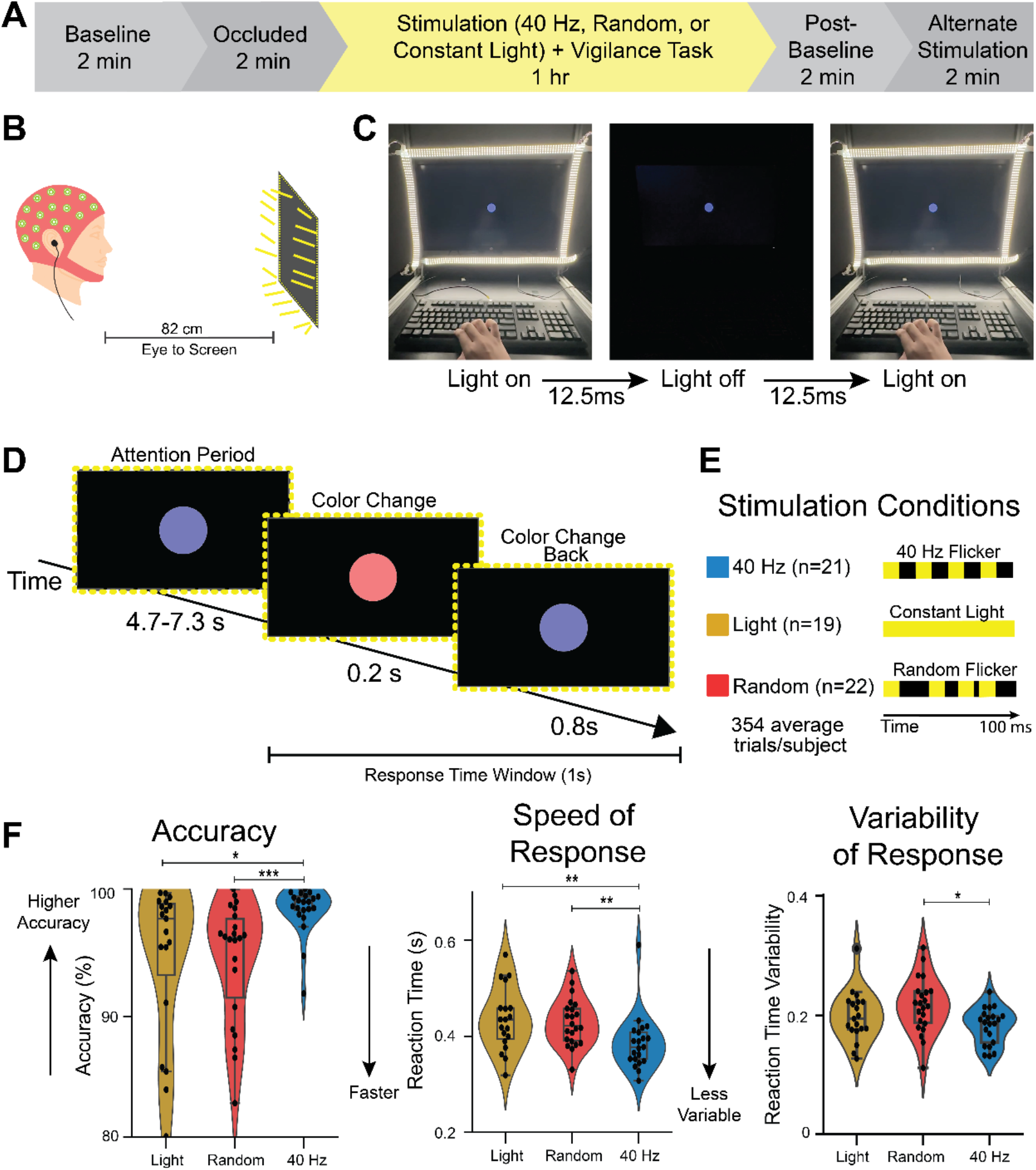
40 Hz flicker improved accuracy and reaction time in a vigilance task. **(A)** Experimental timeline. EEG data were recorded from participants in a baseline period, occluded stimulation period, and during sensory exposure and task performance. **(B)** Illustration of a person undergoing audio and visual stimulation while performing the computer-based psychomotor vigilance task. **(C)** For the 40 Hz flicker group, LEDs on the frame of the monitor turn on and off every 12.5 milliseconds during the task. **(D)** In the task, subjects looked at a blue dot on the screen (“Attending”). At an unpredictable time, the dot changed to red for 0.2 seconds (“Color Change”). After the 0.2 seconds ended, the color changed back to blue (“Color Change Back”). If the subject detected the color change, they pressed a key on a keyboard as quickly as possible and within 1 second of the color change (“Response Time Window”). This schematic shows 1 trial. Subjects performed on average 354 +/-6.76 trials over 1 hour. **(E)** Subjects either underwent 40 Hz flicker (n=21 subjects, blue) in which the LED frame light turned on and off at 40 Hz (on for 12.5 msec and off for 12.5 msec), a control of constant light (n=19 subjects, gold) in which the light stayed on throughout the test, or random flicker (n=22 subjects, red) as a sham condition in which the light turned on for 12.5 msec but turned off for randomized intervals. The random sham stimulation controls for effects of a flickering stimulus, like salience. **(F)** *Left:* Distribution of accuracy for each participant per stimulation group showed that 40 Hz flicker (blue) improved accuracy in the vigilance task compared to both Light (gold, control; p=0.018, ranksum test; q < 0.05, FDR correction for 2 comparisons) and Random flicker (red, sham; p=0.00056, ranksum test; q < 0.05, FDR correction for 2 comparisons). The violin plot shows the histogram of the full distribution with the central box plot spanning the 25^th^ percentile to the 75^th^ percentile of the data. The line inside the box indicates the median, and the whiskers extend to the smallest and largest values within 1.5 times the interquartile range. Each black dot is one subject. *Center:* Distribution of reaction times for each stimulation group shows 40 Hz flicker reduced reaction time in the vigilance task compared to both Light (control; p=0. 0051, ranksum test; q < 0.05, FDR correction for 2 comparisons) and Random flicker (sham; p=0. 0028, ranksum test; q < 0.05, FDR correction for 2 comparisons). *Right:* Distribution of average reaction time variability shows 40 Hz flicker reduced variability in the vigilance task compared to both Light (control, p = 0.261, ranksum test) and Random flicker (sham, p = 0. 0065, ranksum test; q < 0.05, FDR correction for 2 comparisons). Asterisks indicate: *p<0.05, **p<0.01, ***p<0.001

### Study Design and EEG Recording

EEGs were recorded using a 32-channel BioSemi Active Two system with data acquired at a sampling rate of 2048 Hz. In addition to the 32-channel electrode cap, 6 facial electrodes were used including two placed on the mastoids. During each recording, all room lights were turned off, except for the frame of LEDs during the one-hour stimulation portion. The door was closed to shield the participant from light and noise from outside the room. Participants were instructed not to excessively blink, move around unnecessarily, or cross their legs during the study to minimize noise due to muscle movements. EEG data were synchronized to behavior data using TTL parallel port event triggers delivered to the EEG system from the PsychoPy software.

In the study, subjects first underwent a baseline recording followed by an “occluded” recording designed to detect electrical artifacts of stimulation. During the occluded recording, participants wore an eye mask and earplugs while flicker stimulation ran to ensure that detected changes in EEG power in recordings were a result of changes in brain activity rather than a byproduct of electrical noise from stimulation devices. During the one-hour stimulation period, subjects performed a psychomotor vigilance task (PVT, described below).

To be included in analysis, subjects were required to meet criteria showing that 40 Hz audiovisual flicker increased 40 Hz neural activity in some EEG channels, in line with previous studies in humans^10,23,38^. To assess whether subjects exhibited such increases in 40 Hz activity in response to 40 Hz flicker, participants who were not in the 40 Hz group were briefly shown 40 Hz stimulation after the end of their one-hour control or sham stimulation EEG session. For each EEG channel, 40 Hz activity was considered significantly increased in response to flicker if power at 40 Hz exceeded the average power in nearby frequencies (31–39 and 41–49 Hz) by at least three standard deviations. For an individual subject, significant modulation by 40 Hz was defined by the presence of at least three modulated channels, with at least one modulated channel in each hemisphere^23^. Participants who did not exhibit significant 40 Hz modulation were excluded from analysis (n = 2).

### Psychomotor Vigilance Task (PVT)

To assess sustained attention (i.e. vigilance) during stimulation, participants completed a PVT presented via PsychoPy. Participants were instructed to focus on a dot at the center of the screen and respond whenever the dot’s color changed from blue to red (**Fig. 1D**). The “attending period”, or the period when the dot remained blue before the color change, lasted for 4.3-7.3 seconds with the duration jittered per trial to be unpredictable. Then the dot changed colors from blue to red for 0.2 seconds, before switching back to blue for 0.8 seconds, after which a new trial began. During the one hour of the PVT and EEG recording, participants were given short breaks from the task approximately every 20 minutes, during which they remained at the monitor and continued receiving sensory exposure (**Fig. 1E**).

### Behavior Analyses

Each PVT trial was categorized as: (1) a premature response if the participant responded before the color change occurred; (2) a hit if the participant responded within one second after the color change occurred; or (3) a miss if the participant did not respond within that one-second window. Accuracy was computed as the number of hit trials out of total trials and reaction time as the time from color change to a button press. No reaction time was recorded for missed trials or premature responses. Participants with an accuracy below 80% were considered outliers (four in total) and excluded from behavior analyses. This ensured that only participants who were actively engaged in the task and thus continuously receiving stimulation were included in the behavior analysis.

### EEG Data Analyses

Prior to analysis, EEG data were preprocessed to remove noise artifacts such as eye blinks and muscle movements, identified using independent component analysis (ICA), and faulty channels using the EEGLAB toolbox for MATLAB. A high-pass filter at 1 Hz was applied. Data was segmented into epochs, one epoch per trial. Each epoch spanned from four seconds to zero seconds prior to the color change.

To measure the distribution of power contained within the EEG signal over frequency for a given EEG channel, power spectra were calculated using Welch’s method, with 50% overlapping Hamming windows of 2 seconds length and Fast Fourier Transform (FFT) size of 1024, implemented with MATLAB’s Signal Processing Toolbox. We measured normalized power, dividing raw power by the area under the power spectral density from 2-55 Hz to control for individual differences in overall power. We found similar results when normalizing the power spectral density from 1-55 Hz and measuring delta from 1-4Hz.

For visualization of EEG power on scalp maps, FOOOF (fitting oscillations & one over f) was applied to remove aperiodic spectra (https://fooof-tools.github.io/fooof/).

We measured functional connectivity using weighted phase lag index (WPLI) to minimize the impact of volume conduction of source activity^39^. The WPLI varies between 0 and 1 with 0 indicating either no phase coupling or a balanced distribution of leading and lagging phases while 1 denotes the phase of one signal consistently leading the other. WPLI quantifies the imaginary component of coherence using the following formula^39^:

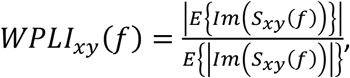

where *S*_*xy*_(*f*)is the cross spectrum between signals recorded from two EEG channels, *x*(*t*)and *y*(*t*)as a function of frequency *f*, where *E*{·}represent cross trial average, *Im*(·)is the Imaginary component. WPLI was computed for every pair of channels within each subject for all attention period epochs.

To identify EEG channel pairs with significant functional connectivity, we compared obtained connectivity values to a null distribution created using a permutation-based approach. In each permutation, we randomly selected an EEG channel from (not necessarily the same channel in each subject) two subjects, then computed the WPLI for the resulting channel pair. This process was repeated 10,000 times to generate a null distribution of connectivity values as we would not expect genuine connectivity across subjects. Real WPLI values were then compared to this null distribution, with WPLI values exceeding the 99.99^th^ percentile (p<0.0001) considered significant. We quantified the total number of significant channel pairs for each frequency band and for each group to identify which bands were significantly modulated by flicker exposure.

### Statistical Approach

In general, a false discovery rate of 0.05 was applied for multiple comparisons. Group comparisons of behavioral measures were assessed using the ranksum test corrected for two comparisons measures (40 Hz versus Random and 40 Hz versus Light). Group comparisons of EEG power were assessed using unpaired, two-sided t-tests corrected for 30 comparisons (2 group comparisons: 40 Hz versus Random and 40 Hz versus Light, 3 channels, and 5 frequency bands). Spearman’s rho values were calculated from correlations between EEG power or WPLI functional connectivity and behavior measures and FDR-corrected for 15 comparisons (3 channels, 5 frequency bands). We identified significant functional connectivity between channel pairs by comparing real WPLI values with shuffled data. Via the permutation test, we generated a threshold that indicated p < 0.0001 (WPLI = 0.089). To identify differences in the proportion of channel pairs with high functional connectivity, measured via WPLI, or with high correlation between functional connectivity and behavior, a chi-squared test was used controlling for two comparisons (40 Hz versus Random and 40 Hz versus Light).

## RESULTS

### 40 Hz flicker improved accuracy and reaction time in a vigilance task

Internally generated gamma oscillations are thought to enhance circuit computations for attention^15,21,40,41^, though the effects of sensory-induced gamma on attention are unknown. Accordingly, we asked how 40 Hz flicker affects accuracy and reaction time in an attentional vigilance task in which subjects must attend to a stimulus over time to detect and respond to an unpredictable color change. To address this question, we exposed subjects to 40 Hz sensory flicker stimulation (40 Hz), constant light (Light) as a control, or random flicker stimulation (Random) as a sham stimulation during a psychomotor vigilance task (PVT) in which participants watching a dot on a screen were instructed to press a key in response to the dot changing colors. Participants with accuracy below 80% were considered not engaged in this simple task and were excluded from the behavior analysis. We compared the accuracy, reaction time, and reaction time variability between the 40 Hz, Light, and Random stimulation groups. Constant light served as a control to see how 40 Hz affects vigilance compared to a steady lighting environment and random flicker was used as sham stimulation to control for the effects that flashing lights and sounds may cause irrespective of their flickering frequency. We found subjects undergoing 40 Hz flicker had faster and more accurate responses in a vigilance task than control and sham stimulation groups. Subjects in the 40 Hz group had a significantly higher accuracy than those in the Light and Random groups during the PVT (**Fig. 1F Left**; 40 vs. Light: p=0.018; 40 vs. Random: p=0.00056, ranksum test; q < 0.05 FDR correction for 2 comparisons; mean accuracy was 98.4%, 94.7%, and 94.5% for 40 Hz, Light, and Random groups, respectively).. Surprisingly, the 40 Hz group also reacted more quickly than the other two groups (**Fig. 1F, middle**; 40 vs Light: p=0.0051; 40 vs. Random: p=0.0028; ranksum test, q < 0.05, FDR correction for 2 comparisons). Reaction times during 40 Hz flicker were on average 11.3% faster than during constant light and 10.0% faster than during Random. Importantly, a speed-accuracy tradeoff^42^, or a higher accuracy correlated with slower reaction time, was not observed (**Supp. Fig 1**). Moreover, there was less variability in the reaction time of 40 Hz-exposed participants than in the Random stimulation group (**Fig. 1F, right**; 40 vs Light: p=0.261; 40 vs.

Random: p=0.0065, ranksum test, q < 0.05, FDR correction for 2 comparisons). Less reaction time variability has been associated with better cognitive performance in prior studies^43–45^. These findings show that 40 Hz flicker enhances vigilance as indicated by higher accuracy and faster reaction time compared to constant light or sham stimulation.

### 40 Hz flicker is correlated with decreases in delta activity during a vigilance task

Because participants undergoing 40 Hz flicker performed better on a vigilance task than control and sham stimulation groups, we wondered how 40 Hz flicker affected multiple oscillation bands that are implicated in attention processing. As described above, endogenous gamma oscillations increase during attention tasks^15,21^. In contrast, elevated delta oscillations (1–4 Hz) have been linked to reduced vigilance, cognitive disengagement, fatigue, attention deficits, increased cognitive workload, decreased alertness, and transitions toward sleep-like states^46–49^. Higher power in beta oscillations (13-30 Hz) is thought to play a role in motor control, sustained attention, and visual attention^50,51^ while higher power in alpha oscillations (8-13 Hz) has been linked to inhibition including the suppression of irrelevant sensory input^29,52^. To determine how these oscillations associated with attention differ between groups, we recorded neural activity using a 32-channel scalp EEG on participants in the 40 Hz, Light, and Random groups (**Fig. 2A**) while they performed the vigilance task during stimulation as described above. To identify neural oscillations associated with attention, we examined neural activity during the attention period preceding the color change of the dot on Hit trials. To perform an unbiased analysis across multiple frequency bands, we measured peak power within five frequency bands: Delta (2-4 Hz), Theta (4-8 Hz), Alpha (8-13 Hz), Beta (13-30 Hz), and Low gamma (30-37 Hz) as well as around the flicker frequency (39-41 Hz). We measured normalized power to control for individual differences in overall power and normalized from 2-55 Hz to minimize the effects of differences in low frequencies on the normalization. We repeated analyses normalized from 1-50Hz and found similar results. We separately measured power around 40 Hz (39-41 Hz, Gamma Flicker Response) as rhythmic sensory stimuli are known to induce a steady state evoked potential^10^. Consistent with prior studies showing periodic flicker increases power in neural activity at the flicker frequency, we found that participants in the 40 Hz group had elevated power in the Gamma-Flicker Response band (**Fig. 2B, C**; 40 Hz vs. Light: Fp1 p=0.00017, Cz p=0.00048, Oz p=0.00095, **Fig. 2B, Supp. Fig. 2**, 40 Hz vs. Random: Fp1 p=7.50E-05, Cz p=0.0033, Oz p=7.06E-05; not corrected for multiple comparisons; unpaired two-sided t-test).

**Figure 2.**
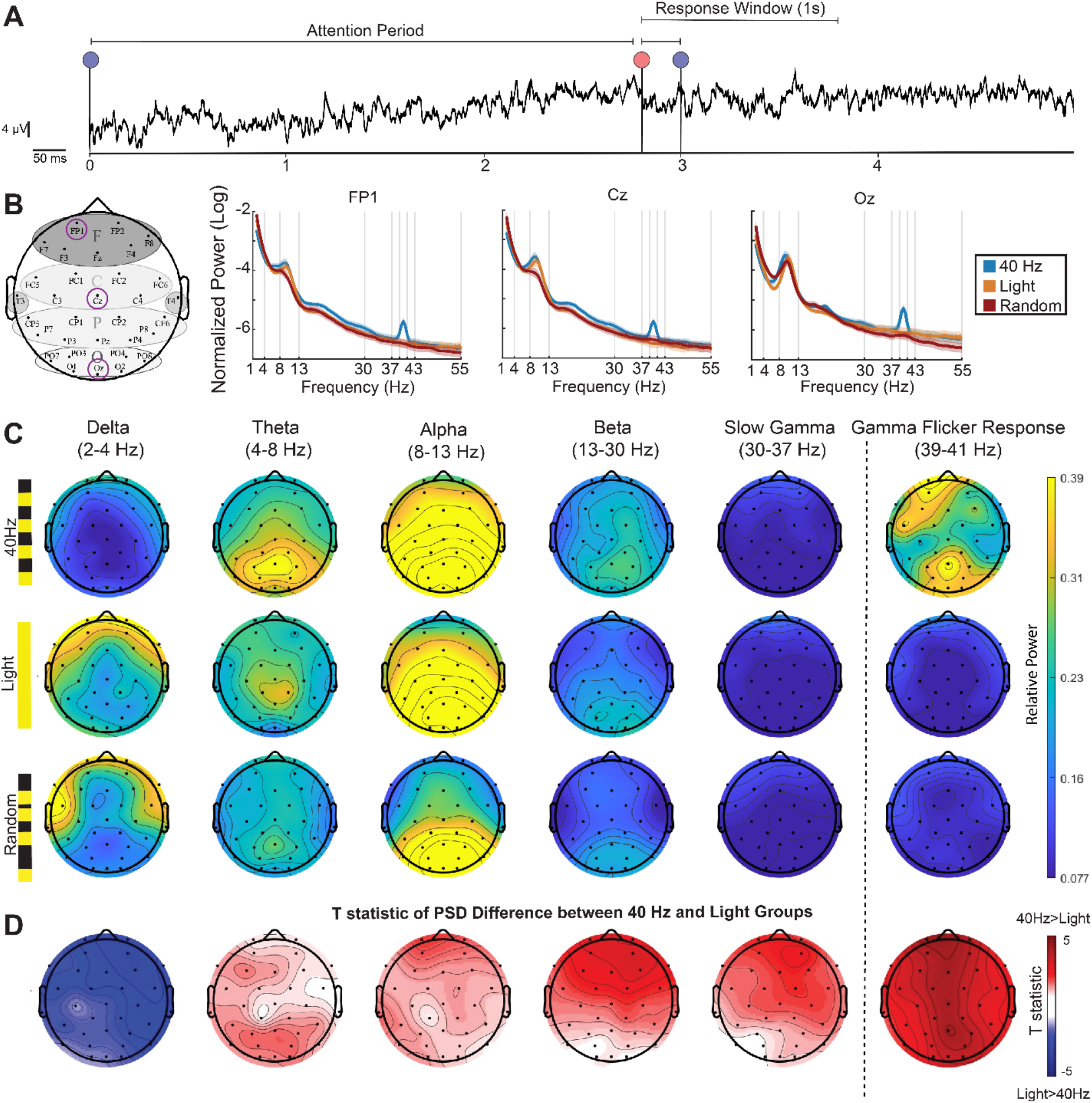
40 Hz flicker decreased delta activity during a vigilance task. **(A)** EEG trace from a single channel during a single trial including the attention period, followed by the color change period, and concluding with the response window. **(B)** *Left:* Topographic map showing the locations of three selected EEG channels (FP1, Cz, Oz). We selected these three channels (Fp1, Cz, and Oz) to capture EEG activity associated with prefrontal/frontal cognitive activity, central sensorimotor processing, and occipital sensory responses. *Right:* corresponding power spectral density (PSD) plots in the 40 Hz (blue), constant light (orange), and Random (red) groups. Each PSD shows mean +/-SEM. **(C)** Topographic maps of periodic components of PSDs extracted using FOOOF (see Methods) for each stimulation group —40 Hz (top row), Light (middle row), and Random (bottom row)— across EEG frequency bands of interest: delta (2–4 Hz), theta (4–8 Hz), alpha (8–13 Hz), slow gamma (30–37 Hz), and the gamma flicker response (centered around 40 Hz). Warmer colors indicate higher power. **(D)** T-statistics of the difference between the real PSD of the 40 Hz group and the Light group during the attention period. Red indicates power in the 40 Hz group is higher than the Light group (positive T-statistics) while blue indicates power in the 40 Hz group is lower than the Light group (negative T-statistics) (two-sided t-test).

We investigated whether there were differences in EEG activity outside the steady state evoked potential expected at 40 Hz. To visualize power across frequency bands in different groups, we used the FOOOF (fitting oscillations & one over f) approach to distinguish rhythmic components of the power spectra from concurrent aperiodic components^53^ (see Methods, **Fig. 2C**). This approach visualizes EEG power above that due to aperiodic changes, largely driven by 1/f modulations, making it easier to compare across frequencies and groups. To assess frontal, central, and posterior EEG activity during vigilance and stimulation, we compared power at Fp1, Cz, and Oz channels between stimulation groups. We selected these three channels (Fp1, Cz, and Oz) to capture EEG activity associated with prefrontal/frontal cognitive activity, central sensorimotor processing, and occipital sensory responses. We were surprised to find lower power within the delta band during 40 Hz flicker compared to the control and sham groups (**Fig. 2B-D**; 40 Hz vs. Light: Fp1 p=0.0064, Cz p=0.016016; **Supp. Fig. 2**, 40 Hz vs. Random: Fp1 p=0.0053, Cz p=0.001, Oz p=0.0055; unpaired t-test; q < 0.05, FDR correction from 30 comparisons from 2 group comparisons, 3 channels and 5 frequency bands; **Table 2**). We did not find significant differences in other frequencies when correcting for multiple comparisons, however we noticed trends of elevated alpha and beta power in the 40 Hz group (**Fig. 2B-D**; beta band 40 Hz vs Light: Fp1 p=0.020; alpha band 40 Hz vs Random Cz: p=0.016, unpaired two-sided t-test; q < 0.05, FDR correction from 30 comparisons from 2 group comparisons, 3 channels and 5 frequency bands; **Table 2**). These results show that 40 Hz sensory flicker not only increases power around the flicker frequency but also decreases delta power during attention.

### Reduced delta band power is correlated with better behavior performance

We then asked how power in these different frequency bands correlated with performance on the psychomotor vigilance task (**Fig. 3A**). We determined if power and behavioral performance were correlated at three key EEG channels of interest: Fp1, Cz, and Oz (**Fig. 3B**). Higher accuracy was significantly correlated with lower delta band power and higher alpha band power (**Fig. 3C**; delta band: Fp1 rho = -0.50, p = 3.2E-05, Cz rho = -0.48, p = 7.1E-05, Oz rho = -0.46, p = 0.00015; alpha band: Fp1 rho = 0.3517, p = 0.0044, Cz rho = 0.3428, p = 0.0056; Spearman’s rank correlation; q < 0.05, FDR correction for 15 comparisons, **Table 3**). Reaction times were positively correlated with increased power in the delta band, meaning lower delta power occurred with faster reaction times. (**Fig 3D**; Delta band: Fp1 rho = 0.4895, p = 5.1E-05, Cz rho = 0.3359, p = 0.0069, Oz rho = 0.3228, p = 0.0096; Spearman’s rank correlation; q < 0.05, FDR correction from 15 comparisons, **Table 3**). Reaction times were negatively correlated with increased power channels within alpha and beta bands meaning higher power in those bands occurred with faster reaction times (**Fig 3D**; Alpha band: Fp1 rho = -0.3337, p = 0.036; Beta band: Fp1 rho = -0.3136, p = 0.012; Spearman’s rank correlation; q < 0.05, FDR correction from 15 comparisons, **Table 3)**. These results show that decreases in delta activity induced by gamma sensory flicker correlated with better vigilance performance. Furthermore, increased alpha and beta activity, which were trended up during gamma flicker, were also correlated with better performance.

**Figure 3.**
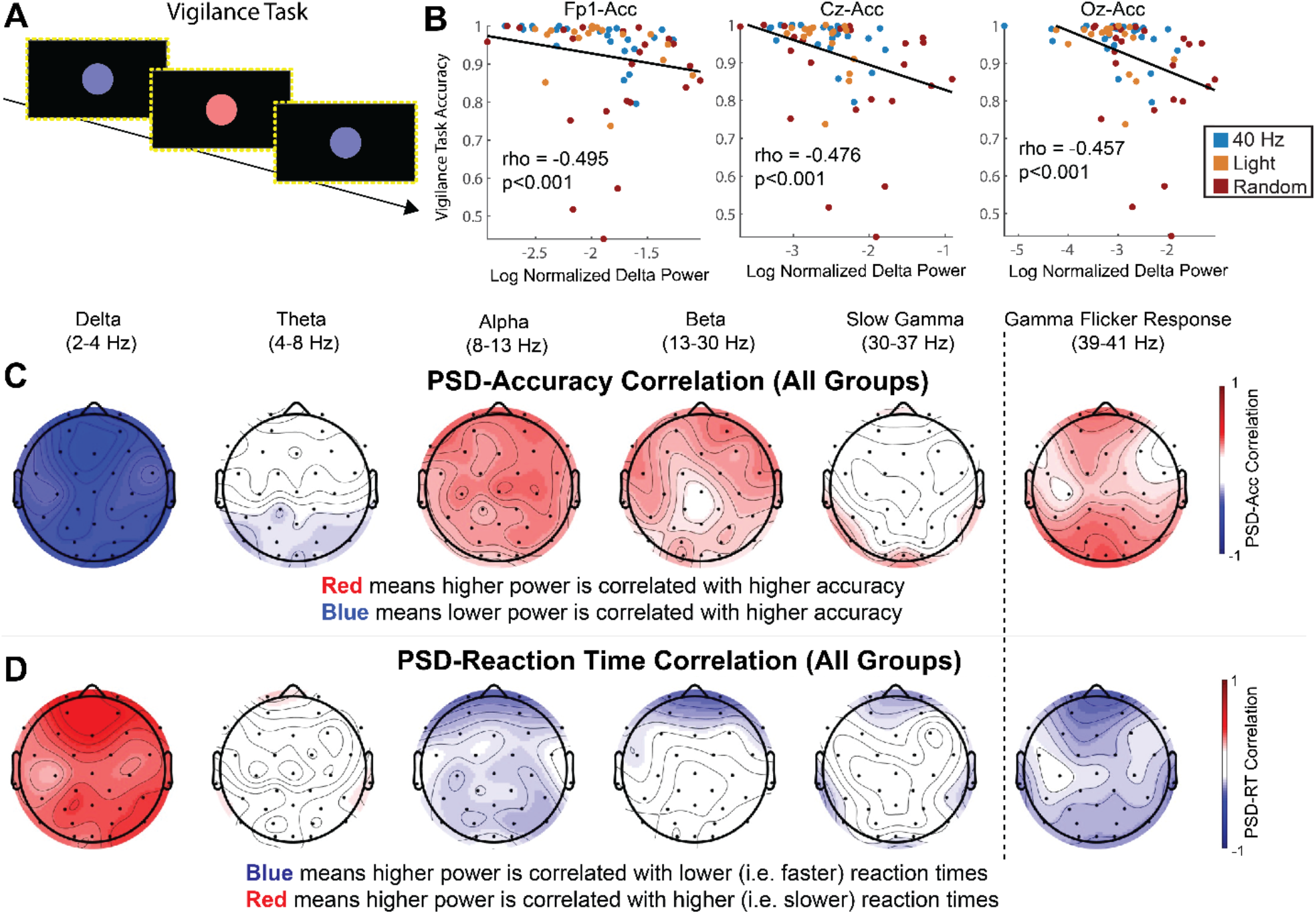
Reduced delta power is correlated with better accuracy and faster reaction time on a psychomotor vigilance task (PVT) **(A)** The sequence of images displayed during the PVT on a monitor with a border of LED lights. **(B)** Spearman’s rank correlation between delta power and accuracy correlation across all subjects on Fp1 (left), Cz (center), and Oz (right). Each dot represents one participant with dot color indicating stimulation group (blue for 40 Hz, gold for Light, and red for Random). **(C)** Power-accuracy correlation topological maps including all experimental groups in the correlation for delta, theta, alpha, beta, slow gamma, and Gamma Flicker Response. Heatmap color shows Spearman’s rank correlation (rho) with red indicating higher power is positively correlated with higher accuracy and blue indicating a negative correlation. **(D)** As in **(C)** for power-reaction time correlation. Heatmap color shows Spearman’s rank correlation with blue indicating a negative power-reaction time correlation, i.e. higher power occurs with faster reaction times or better performance, and red indicating a positive power-reaction time correlation, i.e. higher power occurs with slower reaction times or worse performance.

### 40 Hz flicker increases low-alpha functional connectivity which is correlated with better behavior performance

Because 40 Hz flicker altered oscillations outside the flicker frequency, we then investigated whether it also altered functional connectivity at frequencies outside 40 Hz. High functional connectivity, indicated by correlated fluctuations in neural activity between different channels, is thought to reflect higher communication or coordination between those regions^54,55^. Such coordinated activity is important for cognitive tasks including vigilance^13,16,48,56,57^ (**Fig. 4A**). To assess functional connectivity, we measured the Weighted Phase Lag Index (WPLI) for each pair of EEG channels across each frequency band (**Fig. 4B**).

**Figure 4.**
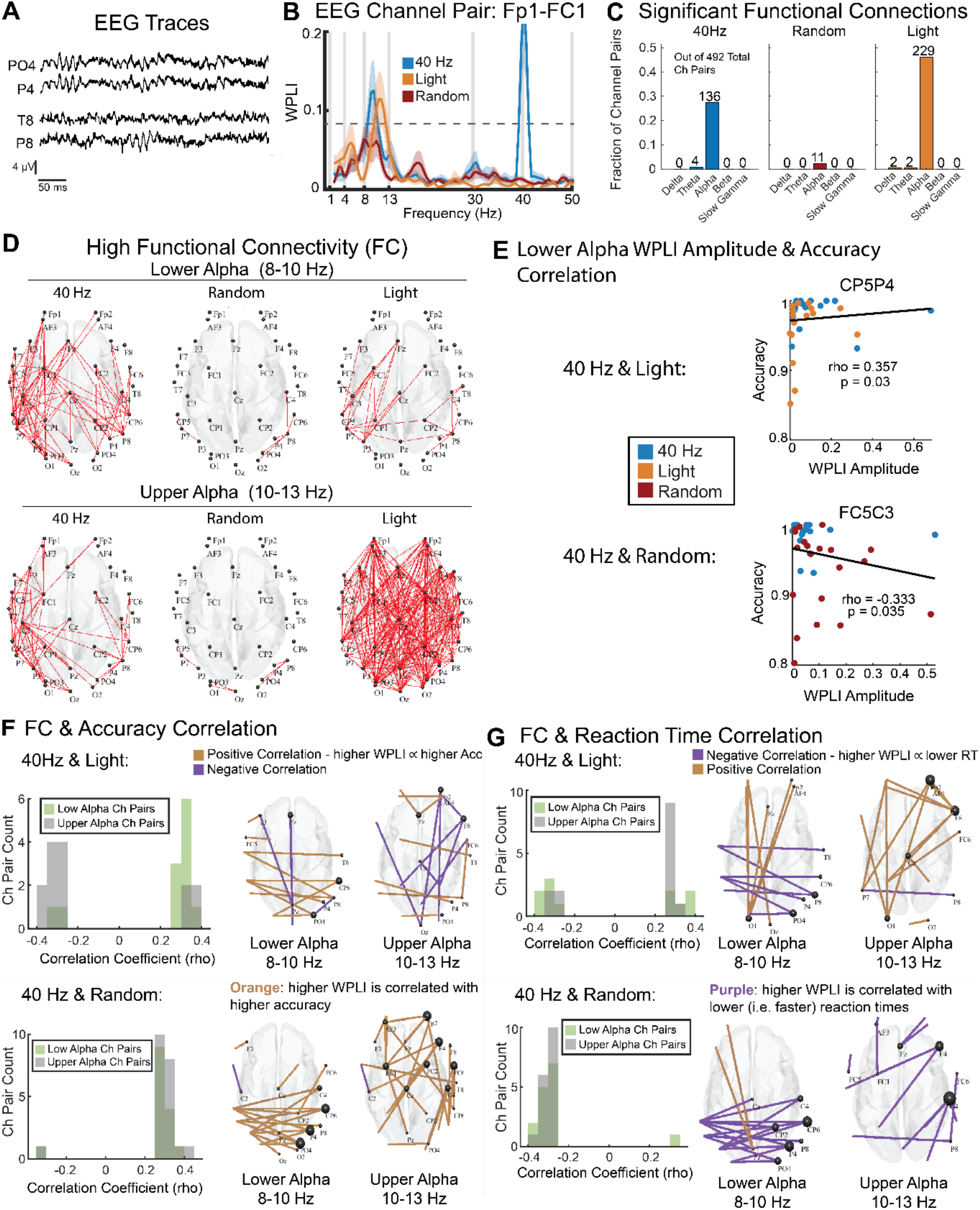
40 Hz flicker increased lower alpha functional connectivity, which correlated with better behavioral performance. **(A)** Representative EEG traces during attention period from two channel pairs with differing functional connectivity. The top panel shows two channels with high functional connectivity, while the bottom panel displays traces from two channels with low functional connectivity as measured via weighted phase lag index (WPLI). **(B)** WPLI as a function of frequency for an example channel pair, Fp1-FC1, across the three experimental groups: 40 Hz flicker (blue), Light (orange), and Random (red). **(C)** Number of EEG channel pairs with significant functional connectivity (p < 0.0001, WPLI > 0.89, permutation test) in each band of interest for each experimental group. **(D)** *Top:* EEG channel pairs (red lines) with high functional connectivity (WPLI values greater than the top quartile of WPLI channel pairs i.e. WPLI > 0.1202) in the lower alpha band (8–10 Hz) in the 40 Hz (left), Random (center), and Light (right) groups. *Bottom:* As above for the upper alpha band (10–13 Hz). **(E)** *Top:* Scatterplot showing a representative EEG channel pair, CP5P4, with a positive correlation between WPLI and accuracy (rho = 0.357, p = 0.03, Spearman’s rank correlation) for 40 Hz and Light groups, indicating that stronger functional connectivity is associated with higher accuracy. Each dot is one subject, with orange for Light and blue for 40 Hz groups. *Bottom:* Scatterplot showing a representative EEG channel pair, FC5C3, with a trending negative correlation between WPLI and accuracy (rho = 0.333, p = 0.035, Spearman’s rank correlation), indicating that stronger functional connectivity is associated with lower accuracy. Each dot is one subject, with red for random and blue for 40 Hz groups. **(F)** *Left:* Distribution of Spearman’s rho-values for channel pairs with trending or significant (p < 0.1) correlation between functional connectivity (WPLI) and accuracy for lower alpha (green) and upper alpha (grey) for 40 Hz and Light (top) and 40 Hz and Random (bottom). In the 40 Hz and Light subjects, strong correlations between lower alpha WPLI and accuracy were more positive, while strong correlations between upper alpha WPLI and accuracy were more negative (top-left, Chi-squared test for proportions, p = 0.005, Cramér’s V: 0.55). In the 40 Hz and random groups, lower and upper alpha functional connectivity were not significantly different (bottom-left, Chi-squared test for proportions, p = 0.81, Cramér’s V: 0.041) and were both positively correlated with accuracy. *Right:* Topological plot of channel pairs with significant WPLI-accuracy correlations for both lower alpha and upper alpha bands. Positive correlations are shown in orange; negative correlations are shown in purple. The channel pairs are displayed for each comparison between groups: 40 Hz and Light (top) and 40 Hz and Random (bottom). **(G)** as in E for reaction time. Note that negative correlations with reaction time indicate correlations with faster performance.

WPLI measures functional connectivity while minimizing the effects that could be due to volume conduction (or correlations at zero-lag). We used this measure to investigate functional networks engaged in this vigilance task while minimizing spurious covariation due to volume conduction. We identified significant functional connectivity between channel pairs by comparing the WPLI threshold of 0.089 from the shuffled data (p < 0.0001, permutation test) within each frequency band of interest (**Fig. 4B**). To determine in which frequency bands’ functional connectivity was prominent, we first quantified the number of channel pairs with significant functional connectivity in each band (**Fig. 4C**). Most of the significant functionally connected (FC) channel pairs (374 out of 384 or 97.4%) were within the alpha band in all groups (**Fig. 4C**). Therefore, we further evaluated how alpha functional connectivity differed between groups.

Examining WPLI values as a function of frequency, we noticed that 40 Hz and Light groups differed in their functional connectivity between lower (8-10Hz) and upper (10-13Hz) alpha bands (**Fig. 4B**). The lower alpha band (sometimes called “low alpha”) is associated with basic sensory and baseline attention processes while the upper alpha band (sometimes called “high alpha”) is more sensitive to memory demands^28^. We found that the peak WPLI frequency within the alpha band was lower in the 40 Hz group (9.35 Hz ± 0.0287 Hz, n = 136 significant FCs) compared to the Light group (9.56 ± 0.0291 Hz, n = 229 FCs; p = 0.000002, ranksum test). Therefore, for further analyses we separated lower and upper alpha. To identify channel pairs with relatively high functional connectivity in each band, we assessed the top quartile of WPLI channel pairs (WPLI>0.12). We found that the 40 Hz group had more channel pairs with high FC in lower alpha than the other groups (**Fig. 4D;** 40 Hz vs Light: X^2^ = 34.06, p < 0.0001, Cramér’s V = 0.1853; 40Hz vs Random: X^2^ = 82.25, p < 0.0001, Cramér’s V = 0.2879; Chi-squared test; q < 0.1, FDR correction from 2 comparisons). In contrast in upper alpha, the 40 Hz group had fewer channel pairs with high FC than the Light group but more channel pairs than the Random group (**Fig. 4D;** 40Hz vs Light: X^2^ = 48.20, p < 0.0001, Cramér’s V = 0.2204; 40 Hz vs Random: X^2^(1) = 188.97, p < 0.0001, Cramér’s V = 0.4365; Chi-squared test; q < 0.1, FDR correction from 2 comparisons). These results show that 40 Hz flicker induced increased FC in more channel pairs within lower alpha than control or sham stimulation.

We then determined whether elevated functional connectivity in lower or upper alpha was correlated with performance in the vigilance task. Because lower alpha is implicated in attention processes, we hypothesized that higher functional connectivity (measured via WPLI) in lower alpha would correlate better with task performance. To see how lower or upper alpha correlated with behavioral performance across subjects and groups, we combined 40 Hz and Random or 40 Hz and Light groups. Because the alpha band FC networks showed different patterns in the 40 Hz, Light, and Random groups, we assessed channel pairs from the 40 Hz and Light groups together and from the 40 Hz and Random groups together to compare correlation patterns between the two group combinations (Fig. 4E). When examining 40 Hz and Light groups, we found lower and upper alpha differed in how functional connectivity was correlated with task accuracy across all channel pairs (**Fig. S4A**; paired t-test, p=0.003). For lower alpha, more channel pairs had significant or trending significant positive correlation (p<0.1) between functional connectivity and accurate performance, in line with our hypothesis (**Fig. 4F** top-left, Chi-squared test for proportions, p = 0.005, Cramér’s V: 0.55). Conversely, when examining 40 Hz and Random groups, we found no significant differences in lower and upper alpha functional connectivity (**Fig. 4F** bottom-left, Chi-squared test for proportions, p = 0.8057, Cramér’s V: 0.040996), and both upper and lower alpha bands were positively correlated with accuracy.

Together, these results show that stronger lower alpha functional connectivity is correlated with more accurate performance in the vigilance task. Thus, the increase in lower alpha functional connectivity in the 40 Hz group is associated with better task performance.

## DISCUSSION

In this study, we found that 40 Hz flicker improved attention in a vigilance task and promoted neurophysiological markers associated with attentional performance. 40 Hz flicker improved accuracy and reaction times in a vigilance task compared to both sham stimulation and constant light, indicating heightened attentional performance. Looking at scalp electrophysiology, we observed expected increases in gamma power, aligning with prior studies on gamma flicker^6,10,23^. Unexpectedly, we also found that 40 Hz flicker decreased delta power during attention periods and found trends of elevated alpha and beta power. Importantly, decreased delta power was correlated with better behavioral performance. Assessing functional connectivity during attention, 40 Hz flicker increased lower alpha functional connectivity, and this increase was correlated with enhanced task performance, suggesting a link between increased lower alpha coherence and improved vigilance. Previous studies have primarily explored the neural health benefits of *chronic* gamma flicker, especially in the context of neurodegenerative diseases^1,3^. In contrast, our study offers novel insights into the *acute* effects of 40 Hz flicker on attention and multi-band oscillatory dynamics in healthy adults. By providing evidence of immediate neurophysiological and behavioral effects, we enhance our understanding of how gamma flicker impacts attention-related processes in healthy adults.

This study broadens the application of gamma flicker from prior work in disease to attentional enhancement in non-clinical contexts. Furthermore, prior flicker studies examined the flicker effects on EEG power at the frequency of flicker while we examined effects at a wide range of frequencies. Surprisingly, our EEG data also showed that 40 Hz subjects had significantly different power in specific neural frequency bands outside the gamma range compared to the control groups. By documenting changes in both power and functional connectivity across multiple frequency bands, we provide novel insights into how exogenously induced gamma oscillations affect other oscillatory bands. Notably, 40 Hz flicker decreased delta and increased lower alpha connectivity, modulations that were correlated with improved behavioral performance. Our results demonstrate the potential for acute 40 Hz flicker to enhance behavioral performance in a vigilance task. This application could be beneficial in jobs where prolonged attention is critical, such as in air traffic control, surgical monitoring, piloting, or long-distance driving. These findings also suggest the potential for flicker-based interventions in non-clinical settings, such as students studying for an exam. However, as our study focused on healthy young adults, additional research is needed to verify these effects in other populations, particularly those with attention deficits.

Our findings pave the way for future studies to further explore gamma flicker’s potential effects on attention processing. Because we measured neural activity using scalp EEG, our findings largely result from cortical activity. Further studies could investigate the effects of 40 Hz flicker on deep and spatially specific neural structures involved in attention. We identified significantly connected channel pairs in each frequency band, and future work could further dissect how stimulation alters the different spatial patterns of connectivity. We observed the effects of 40 Hz flicker in a single one-hour session. Chronic exposure studies could test if gamma flicker yields cumulative attentional benefits, or greater benefits after repeated exposures, further supporting its application in cognitive enhancement tools or therapeutic interventions for both healthy and clinical populations. The acute impacts observed here in healthy adults may differ in other demographics, such as clinical populations (e.g., Alzheimer’s, ADHD, epilepsy, etc.) or older adults, who may not respond to gamma flicker in the same way as younger individuals^1,3,58,59^. Future research could explore whether different types of stimulation—such as audio-only, tactile, invisible spectral^4,60^, transcranial electrical or magnetic stimulation, or 40 Hz-modulated multimedia—and other stimulation frequencies produce comparable effects.

This study bridges an important gap between therapeutic gamma flicker research in clinical populations and cognitive enhancement in healthy populations. Our findings of immediate attentional improvement highlight gamma flicker’s potential as a non-invasive tool for cognitive enhancement and suggest applications for attention-demanding contexts. The observed multi-band oscillatory changes extend the current understanding of gamma flicker’s role, offering evidence that exogenous gamma oscillations may serve distinct and complementary functions to endogenously generated gamma rhythms. By identifying acute behavioral benefits and neural changes of gamma flicker, this research supports a wider range of applications and encourages further investigation into the mechanisms and utility of gamma oscillation entrainment across cognitive domains.

## Acknowledgements

We thank the Singer lab for valuable comments on the manuscript and technical assistance. We thank Vanesa Vargas, Shangze Lyu, Natalie Black, Luke Braun, Yan Xie, Sophia Baker, William Dunne, Sadat Uddin, Michael Waleign, Anna Zagora, Ben Borron, Victor Nguyen, Lyndah Lovell; Qiliang He; Vishwadeep Ahluwalia and the Center for Advanced Brain Imaging (CABI), Emily Hokett, Kyoungeun Lee, and Tiffany Nguyen for technical assistance.

## Author contributions

MKA: conceptualization, methodology (stimulation during EEG and behavior assays), software, investigation, data curation, analysis, visualization, writing – original draft, writing – review and editing. LZ: conceptualization, methodology (measuring functional connectivity), software, visualization, supervision, writing – original draft, writing – review and editing. SM: software, investigation, data curation, analysis, writing – review and editing. ACS: conceptualization, writing – original draft, writing – review and editing, supervision, project administration, funding acquisition.

## Ethics

The study was approved by the Institutional Review Board at the Georgia Institute of Technology and was conducted according to the principles expressed in the Declaration of Helsinki.

## Data and code availability

Data from this study will be made available on OpenNeuro upon publication: https://doi.org/10.18112/openneuro.ds006222.v1.0.0. Code is available on GitHub: https://github.com/singerlabgt/FlickerEEGAttention_HealthyAdults_Manuscript.

