## Supplementary Information for "40 Hz Audiovisual Stimulation Improves Sustained Attention and Related Brain Oscillations"

Contents:
3 Tables
4 Figures

**TABLES**

**Table 1. Demographics of Stimulation Subjects per Group**

| Characteristics | 40Hz | Random | Light | 40Hz vs Random p-value | 40Hz vs Constant p-value |
|---|---|---|---|---|---|
| | (n = 21) | (n = 22) | (n = 19) | | |
| Years of Age (mean, (s.d)) | 20 (1.4) | 20 (1.6) | 21 (4.0) | 0.42 | 0.29 |
| Female Sex (n (%)) | 7 (33) | 13 (59) | 10 (52) | 0.13 | 0.61 |
| Years of Education (mean, (s.d)) | 14 (1.7) | 15 (1.5) | 16 (2.3) | 0.38 | 0.12 |
| Race: Caucasian (n (%)) | 9 (43) | 10 (45) | 8 (42) | 0.99 | 0.88 |

Years of Age, Years of Education, and all Percentage values are rounded to the nearest integer. Standard deviation values are rounded to the nearest tenth. Student T-tests were used to compare Years of Age and Years of Education. Chi-square tests were used to compare Female Sex, and Race. P-values were not corrected for multiple comparisons.

**Table 2. Difference between 40 Hz & Light Group and 40 Hz & Random Groups**

| **Band** | **Channel** | **40 vs Light** | **40 vs Random** |
|---|---|---|---|
| **Delta** | Fp1 | 0.048 | 0.048 |
| **Delta** | Cz | 0.0805 | 0.031 |
| **Delta** | Oz | 0.3931 | 0.048 |
| **Theta** | Fp1 | 0.5278 | 0.2967 |
| **Theta** | Cz | 0.7102 | 0.8719 |
| **Theta** | Oz | 0.5614 | 0.6218 |
| **Alpha** | Fp1 | 0.2967 | 0.2967 |
| **Alpha** | Cz | 0.5567 | 0.0805 |
| **Alpha** | Oz | 0.5567 | 0.5565 |
| **Beta** | Fp1 | 0.0855 | 0.1416 |
| **Beta** | Cz | 0.4385 | 0.2863 |
| **Beta** | Oz | 0.8719 | 0.6098 |
| **Slow Gamma** | Fp1 | 0.2967 | 0.2863 |
| **Slow Gamma** | Cz | 0.3786 | 0.4164 |
| **Slow Gamma** | Oz | 0.6218 | 0.2967 |

T-test comparing the power between 40 Hz and Light groups and 40 Hz and Random groups in each frequency band on the channels of interest. P-values are FDR-corrected for 30 comparisons (3 channels, 5 frequency bands, 2 comparisons).

**Table 3. Power-Behavior Correlation**

| | | PSD-Acc | | PSD-RT | |
|---|---|---|---|---|---|
| **Channel** | **Band** | **rho** | **p-value** | **rho** | **p-value** |
| Fp1 | Delta | -0.495 | 0.0005 | 0.4895 | 0.0008 |
| Cz | Delta | -0.4759 | 0.0005 | 0.3359 | 0.0358 |
| Oz | Delta | -0.457 | 0.0007 | 0.3228 | 0.0358 |
| Fp1 | Theta | -0.1421 | 0.3283 | 0.1943 | 0.232 |
| Cz | Theta | -0.0765 | 0.5874 | -0.0077 | 0.9958 |
| Oz | Theta | -0.2226 | 0.1284 | 0.1079 | 0.539 |
| Fp1 | Alpha | 0.3517 | 0.0164 | -0.3337 | 0.0358 |
| Cz | Alpha | 0.3428 | 0.0167 | -0.2108 | 0.2119 |
| Oz | Alpha | 0.2528 | 0.084 | -0.1815 | 0.2513 |
| Fp1 | Beta | 0.2517 | 0.084 | -0.3136 | 0.0358 |
| Cz | Beta | 0.1199 | 0.3982 | 0.0277 | 0.9958 |
| Oz | Beta | 0.1518 | 0.315 | -0.0021 | 0.9958 |
| Fp1 | Slow Gamma | 0.1607 | 0.3068 | -0.2081 | 0.2119 |
| Cz | Slow Gamma | 0.0457 | 0.72 | 0.0007 | 0.9958 |
| Oz | Slow Gamma | 0.2583 | 0.084 | -0.1591 | 0.313 |

Spearman's rank correlation rho-values and p-values for correlations between power and accuracy and between power and reaction time.  p-values are FDR-corrected for 15 comparisons (3 channels, 5 frequency bands).

### SUPPLEMENTARY FIGURES

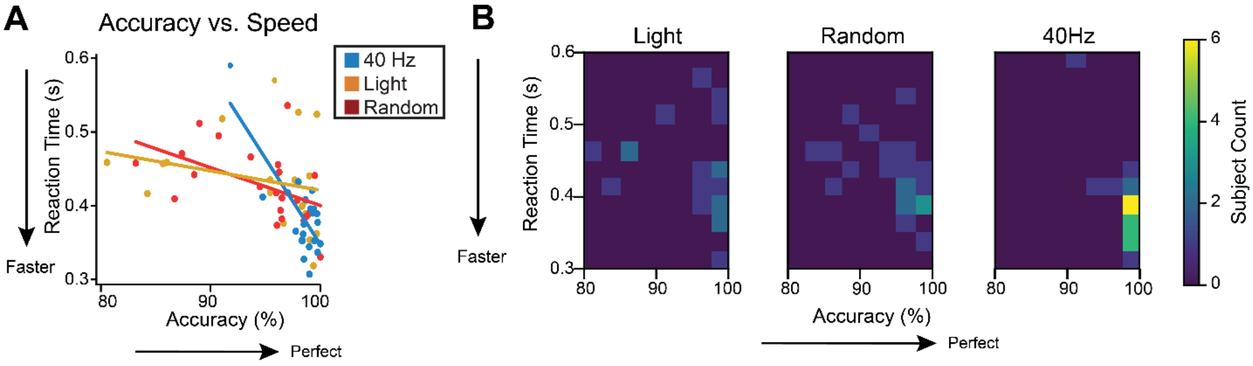


**Supplementary Figure 1. Accuracy versus speed relationship during PVT**

**(A)** Average accuracy versus average reaction time per subject for those that underwent 40 Hz flicker (blue; Spearman's rank correlation: Spearman's rho=-0.412, p-value= 0.072, n=21), Random (red; Spearman's rank correlation: Spearman's rho=-0.569, p-value=0.006, n=22), and Constant Light (gold; Spearman's rank correlation: Spearman's rho=-0.412, p-value=0.079, n=19). Each dot is one subject. Lines indicate best linear fit (first-degree polynomial fit). Higher accuracy was not correlated with slower reaction times. Instead, there was a trend of a correlation (40Hz and light groups) or a significant correlation (Random group) in the opposite direction (faster reaction time was correlated with higher accuracy).

**(B)** Heatmap of average accuracy versus average reaction time per stimulation group with more yellow colors indicating more subjects in that bin.

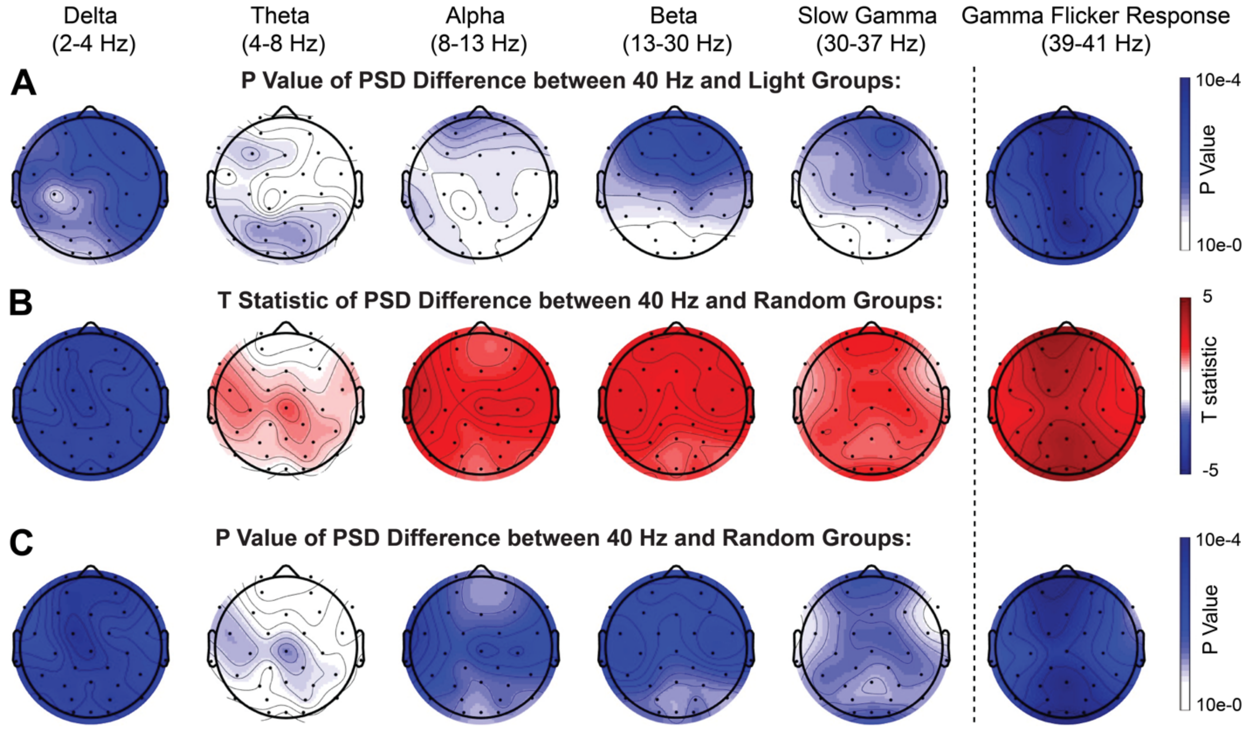


**Supplementary Figure 2. 40 Hz flicker decreases delta activity during a vigilance task**
**(A)** maps of p-values (not corrected for multiple comparisons) for PSD differences from comparing the PSD of the 40 Hz group and Light groups during the attention period.
**(B)** PSD difference map showing the T statistic of the difference between the PSD of the 40 Hz group and Random groups during the attention period.
**(C)** maps of p-values (not corrected for multiple comparisons) for PSD differences from comparing the PSD of the 40 Hz group and Random groups during the attention period.

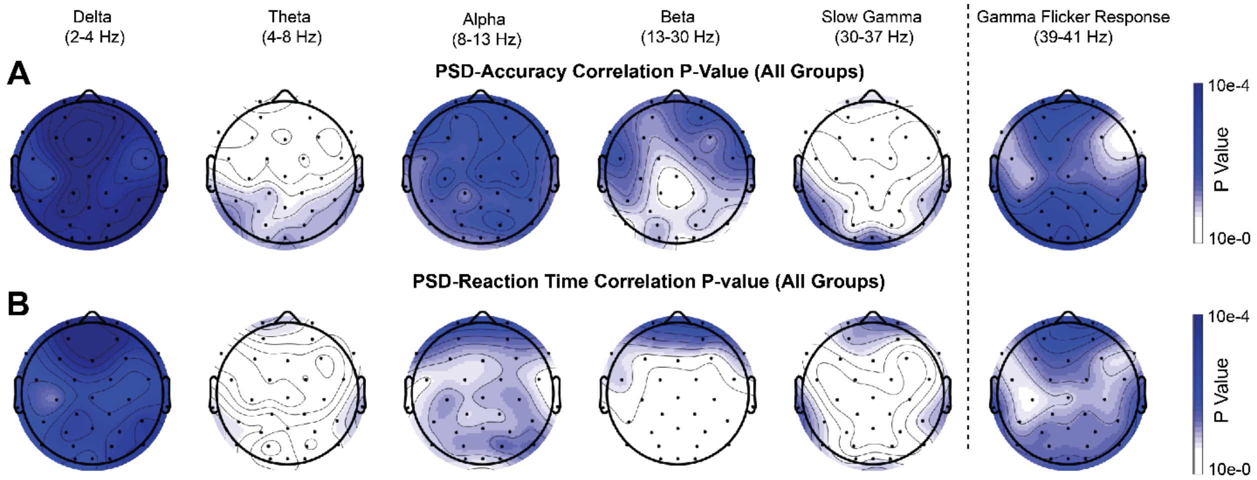


**Supplementary Figure 3. Power-behavior correlation maps of p-values.**

**(A)** Maps of p-values (not corrected for multiple comparisons) for power-accuracy correlation across all subjects in all groups. Spearman's rank correlation.

**(B)** Maps of p-values (not corrected for multiple comparisons) for power-reaction time correlation across all subjects in all groups. Spearman's rank correlation.

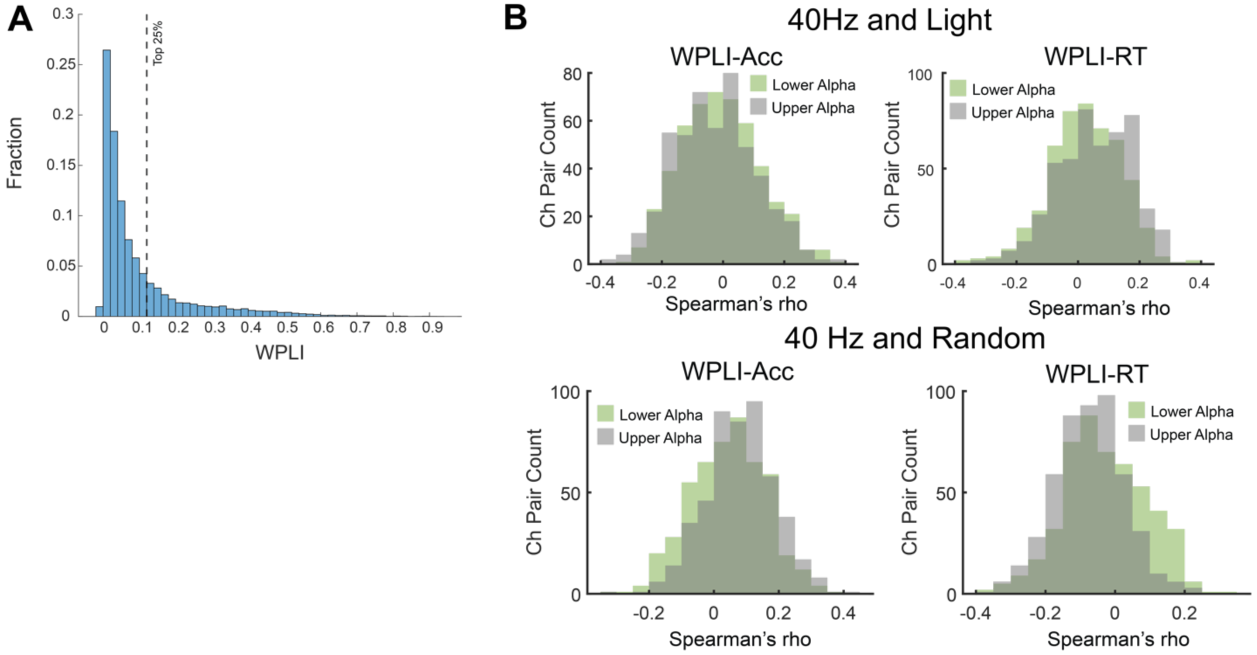


**Supplementary Figure 4. WPLI and WPLI-behavior correlation distributions**

**(A)** Distribution of peak WPLI values within the alpha band of all subjects' channel pairs with top quartile (WPLI = 0.12) indicated by dashed line (n = 67 subjects x 496 channel pairs = 33,232 alpha peak WPLI values).

**(B)** Distribution of WPLI-Behavior Spearman's rank correlation coefficients (rho) across all channel pairs (n=496) comparing lower alpha (green) and upper alpha (grey) frequency bands. *Top left*: Distribution of WPLI-Accuracy Spearman's rho values for the 40 Hz and Light conditions. The distributions of lower alpha and upper alpha were significantly different (paired t-test, p = 0.003). *Top right*: Distribution of WPLI-Reaction Time Spearman's rho values for the 40 Hz and Light conditions. The lower alpha and upper alpha distributions were significantly different (paired t-test, $p = 4 \times 10^{-9}$, n=496). *Bottom left*: As in top left for 40 Hz and Random conditions. The lower alpha and upper alpha distributions were significantly different (paired t-test, $p = 5 \times 10^{-10}$, n=496). *Bottom right*: As in top right for 40 Hz and Random conditions. The lower alpha and upper alpha distributions were significantly different (paired t-test, $p < 10^{-16}$, n=496).
